# Variation in enhancer activity across brain regions defines neurological disease risk and shapes cellular pathology

**DOI:** 10.64898/2026.09.18.752151

**Authors:** Hannah Ramcharan, Yasmine Sami, Kathryn Hazel, Chidera Okeke, Zia Barnard, An T. Hoang, Scott McCallum, Chesna Apere, Wenjuan Du, Megan Madden, Yike Huang, Xochitl Luna, Gesmira Molla, Deborah Mash, Olivia Corradin

## Abstract

**Background:** Genetic variants associated with neurological traits are enriched in enhancers active in neurons. However, there are many types of neurons with varying functions across the brain, making it difficult to pinpoint insights into disease pathology. We set out to investigate whether cell type and brain region specificity of transcriptional enhancers could reveal new insights into the cellular pathology of neurological traits.

**Results:** We performed H3K27ac ChIP-seq on neurons and glia sorted from post-mortem tissue for six brain regions in triplicate and integrated these datasets with existing single-cell ATAC-seq data. While 87% of neuronal enhancers had similar activity across brain regions, we identified 40,049 neuronal regulatory elements that vary in activity across brain regions, which we termed Brain Region Variable Elements (BRVEs). Some BRVEs reflect differences in cell composition, such as a high proportion of medium spiny neurons (MSNs) in the nucleus accumbens, whereas others capture developmental trajectories, including enhancers with robust activity across telencephalic regions that are inactive in the diencephalic hypothalamus. Genetic variants within BRVEs are disproportionately enriched for the heritability of neurological traits, capturing far more heritability than neuronal enhancers with uniform activity across the brain. Additionally, genes linked to BRVEs are also more likely to be implicated in neuropsychiatric disorders by differential expression studies. Finally, we present a new method, “GWAS-LOCATE” that leverages variation in enhancer activity across the brain to assign 23,494 GWAS loci to specific cell types and brain regions, including 9.8% linked to MSNs. This includes a BMI risk locus linked to the transcription factor ISL1. ISL1 knockdown in iPSC-derived MSNs revealed differentially expressed genes enriched for BMI heritability, supporting a role for ISL1-mediated MSN pathways with BMI.

**Conclusions:** Collectively, these findings demonstrate that the enhancer specificity across the brain provides a powerful framework for dissecting complex trait biology and revealing cellular pathology.

## Introduction

Transcriptional enhancer elements are noncoding genomic regions that play a critical role in regulating gene transcription. Enhancers are highly cell type-specific and dynamically responsive to changes in the cellular environment, enabling precise spatiotemporal control of gene expression. Disruption of cis-regulatory elements such as enhancers is thought to be a major contributor to human disease, accounting for an estimated 80% of known genetic risk [1]. Disease-associated genetic variants are enriched in enhancers that are active in cell types relevant to disease pathology [2–4]. For example, variants associated with neurological and neuropsychiatric disorders map to enhancers active in brain tissues [5–7]. However, such genome-wide enrichment analyses, while informative about the main contributing cell types, do not pinpoint the specific cell type or cellular context affected by individual genetic risk factors. Moreover, there are many neuronal subtypes with distinct functions across brain regions [8, 9]. Further studies are therefore needed to link genetic variants to specific neuronal subtypes and brain regions to better understand the cellular pathology of neurological traits.

Distinct brain regions have long been associated with specialized functions. Early lesion studies have shown that damage to specific areas leads to characteristic behavioral and functional deficits. Additional studies have corroborated these findings through neuroimaging, in which specific functions are linked to regional network activity through neurostimulation [10, 11]. Post-mortem brain tissue studies of neurological disease have profiled differential gene expression patterns across multiple brain regions for a given disease. These studies often reveal distinct sets of differentially expressed genes (DEGs) for each evaluated brain region. Some of these differences are derived from cell type-specific genes such as DEGs identified in the nucleus accumbens that are more robustly expressed in GABAergic medium spiny neurons (MSNs). In other cases, DEGs are robustly expressed in multiple brain regions, yet are only differentially expressed in one region [12–15].

Studies leveraging ChIP-seq or open chromatin profiling, such as ATAC-seq, have shown that enhancers active in neurons are particularly enriched for genetic variants associated with neuropsychiatric and neurodegenerative diseases [16, 17]. Enhancers that are specific to neuronal compared to glial cells, have the strongest contributions to disease heritability, with some notable exceptions, such as Alzheimer’s disease, which is enriched in microglia active enhancers [5–7]. Bulk and single-cell profiling of human brain tissues have greatly expanded our understanding of cell composition and diversity within the brain, identifying hundreds of unique cell clusters across many brain regions using transcriptomic, ChIP-seq and ATAC-seq data [5, 7, 8, 18–20]. Here we incorporate H3K27ac ChIP-seq from healthy donors for six brain regions (prefrontal cortex, orbitofrontal cortex, hippocampus, hypothalamus, nucleus accumbens and amygdala) with prior post mortem brain studies to evaluate enhancer activity across the brain. H3K27ac is associated with active enhancers and promoters. H3K27ac is more cell type-specific and has higher correlation with active transcription than open chromatin alone, as ATAC-seq can identify open chromatin regions that are bound by either transcriptional activators or repressors [21–23]. We identified a subset of neuronal cis regulatory elements that vary in activity across the brain (brain region variable elements, BRVEs). These regulatory elements are strongly implicated in human disease, accounting for a disproportionate share of the heritability of neurological traits and exhibiting significant enrichment near genes differentially expressed in neuropsychiatric disorders. We further compare individual GWAS loci with region and cell type-specific regulatory element activity patterns to link individual alleles to their likely cell type of action. While glial-specific elements are not globally enriched for neurological trait heritability, we identify approximately 20% of neurological risk loci predicted to influence glial cells. Thus, while not the dominant cell type for these traits, the risk architecture of most traits includes some glial-acting risk loci. We also highlight the value of this approach in expanding understanding of neuronal subtypes relevant to disease pathogenesis by investigating the role of the ISL1 risk locus in medium spiny neurons for Body Mass Index (BMI).

## Results

### H3K27ac across six brain regions identifies regulatory elements with variable activity patterns across cell type and brain region

We performed ChIP-seq in triplicate for H3K27ac, a marker of active enhancers and promoters, on NeuN+ and NeuN-nuclei isolated from post-mortem tissue of healthy donors across six brain regions: the prefrontal cortex (PFC), orbitofrontal cortex (OFC), hippocampus (Hipp), amygdala (Amyg), nucleus accumbens (NAc), and hypothalamus (Hypo) (Fig 1A, Table S1). H3K27ac peak activity for NeuN+ cells showed high correlation with human *ex vivo* neurons and postmortem neurons, and poor correlation with *ex vivo* microglia, *ex vivo* astrocytes, and postmortem glial cells, confirming stratification of neuronal versus non-neuronal cells in our data (Fig S1A). NeuN+ and NeuN-cell populations also showed expected H3K27ac enrichment at neuronal marker genes TUBB3, GRIN1 and GAD1 and glial marker genes MBP, SLC1A3 and TMEM119, respectively (Fig 1B). We also observed H3K27ac enrichment for broad regional specific markers, such as FOXG1 for telencephalic brain regions, NEUROD6 and SATB2 for cortical regions, MEIS2 for the subpallium and NKX2-1 for the hypothalamus (Fig 1C).

**Figure 1.**
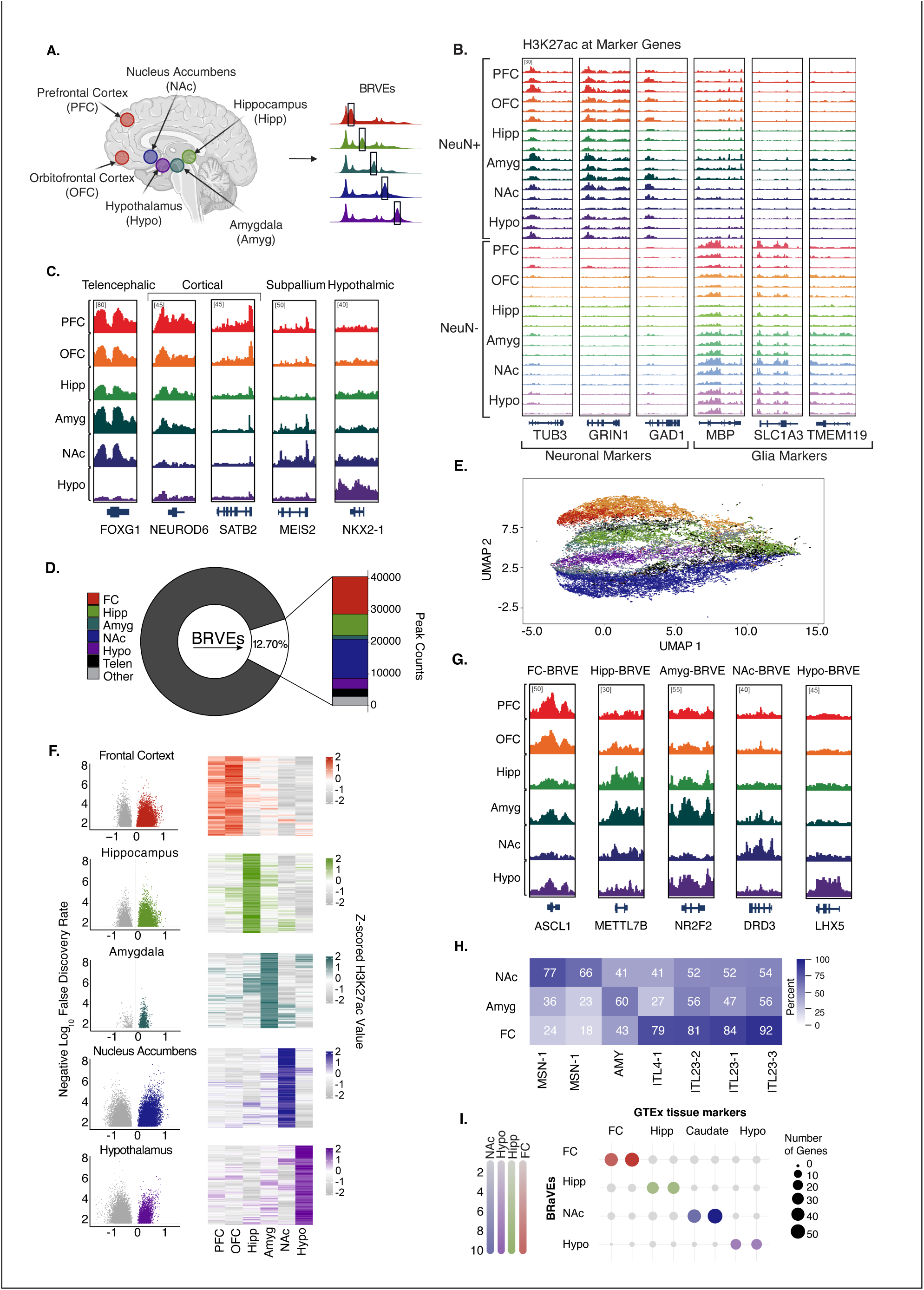
Identification of BRVEs and Across Six Different Brain Regions: **(A)** ChIP-sequencing of H3K27ac was conducted on six brain regions. BRVEs are defined as H3K27ac peaks which showed ANOVA significant variation across these regions. **(B)** H2K27ac activity of canonical neuronal and glial marker genes in NeuN+ and NeuN- cells. **(C)** H2K27ac activity at marker genes for broad regional specific markers for telencephalic, cortical and subpallium hypothalamic regions. **(D)** Donut plot showing the percentage of BRVEs (white) we identified from the total number (n=315,334) of peaks (black). The zoomed in section shows number of BRVEs identified for each brain region including the frontal cortex (FC, red), hippocampus (Hipp, green), amygdala (Amyg, teal), nucleus accumbens (NAc, blue), hypothalamus (Hypo, purple), telencephalic (black) and unclassified BRVEs (light grey). **(E)** UMAP of the BRVEs colored by brain region using color described (D). **(F)** (Left) Volcano plots showing the log₂ fold change in peak signal for each brain region relative to the mean peak signal across the remaining brain regions (x-axis) and the −log₁₀-transformed ANOVA *p*-value (y-axis). Dots in color represent BRVEs. (Right) Heatmaps showing hierarchically clustered peak signals after row-wise z-score normalization across brain regions. **(G)** H2K27ac activity of representative BRVEs overlapping marker genes for each region. The rows represent the H2K27ac ChIP tracks and the columns represent the representative BRVE for each brain region. **(H)** Heatmap showing the percentage of ATAC peaks overlapping BRVEs that are active in each cell type. Percentages were calculated as the number of overlapping ATAC peaks active in a given cell type divided by the total number of brain-region BRVEs overlapping the corresponding brain snATAC peaks. Rows indicate BRVEs and columns represent the snATAC cell types. **(I)** Gene enrichment analysis comparing BRVEs to GTEx V8 tissue signatures using Male [M] (left) and Female [F] (right) ages 50–59 Up gene sets from Frontal Cortex (BA9) (red), Hippocampus (green), Caudate [Basal Ganglia] (blue), Hypothalamus (purple. Dot size reflects the number of genes contributing to each term within a given brain region, and color indicates the −log₁₀ transformed enrichment p-value.

We applied Analysis of Variance (ANOVA) to identify putative cis-regulatory elements that showed variability in H3K27ac across the brain (BRVEs). While 87% of H3K27ac peaks had similar activity across the brain, we identified 40,049 BRVEs (FDR 0.10) and 14,976 BRVEs (FDR 0.05) with significant variation in activity patterns (Fig 1D, left). To classify BRVEs based on the brain region specificity patterns, we ran post-hoc t-tests comparing each brain region to all others for all BRVEs (FDR 0.10). This reveals the subsets of BRVEs that are associated with region-specific activity. Plotting BRVEs in a low-dimensional UMAP embedding revealed distinct clustering according to brain region association (Fig 1E). The OFC and PFC associated BRVEs showed a high degree of similarity, consistent with prior studies highlighting that these brain regions are not only anatomically close but also share similar cell compositions [24]. Thus, we combined PFC and OFC samples in t-tests to define “frontal cortex” associated BRVEs (FC-BRVEs) (Fig 1F). Representative region-associated BRVEs for each region are shown in Figure 1G. NAc- and FC-BRVEs were the largest classes, representing 30.7% and 29.3% of BRVEs, respectively (Fig 1D,F). We compared BRVEs to snATAC datasets for available brain regions and found high concordance in activity and specificity (Fig 1H) [5]. 73% of BRVEs were identified as open chromatin regions in the brain in this dataset. Of the NAc-BRVEs that coincide with ATAC regions, 66–78% were active in MSN clusters, whereas only 40–54% were active in intratelencephalic (IT) cortical neuron clusters. Conversely, among FC-BRVEs, 79–92% were active in IT neuron clusters, whereas only 18–24% were active in MSN clusters. We also found that the nearest gene to BRVEs to have region-specific gene expression patterns showing enrichment in brain region marker gene sets defined by the GTEx consortium (Fig 1I).

Thus, we identified a subset of regulatory elements with varying activity patterns across the brain (BRVEs), which can be classified by region-specific activity (BRVEs), as supported by concordance with snATAC-seq and gene expression patterns.

### BRVE are disproportionately enriched for disease heritability

We next examined genetic variants overlapping BRVEs. Using stratified LD score regression (s-LDSR), we estimated heritability enrichment across 65 traits, including 53 neuropsychiatric and neurodegenerative disorders as well as 12 metabolic and autoimmune traits for comparison. We observed robust heritability enrichment within BRVEs across 29 neurological traits. We compared this enrichment to n-matched active neuronal enhancers that have consistent H3K27ac activity across all brain regions. 16 of the 29 neurological traits that were enriched in BRVEs were not significantly enriched in enhancers with consistent activity patterns across the brain (Fig 2A). Of the remaining 13 traits, 11 had higher heritability enrichment in BRVEs than peaks with consistent activity (p-value = 3.6E-4, Fig 2B,C). Overall, we observed significantly higher per SNP enrichment for BRVEs compared to peaks with consistent activity (p-value = 3.6E-4, Fig. 2C). We did not observe heritability enrichment for the 12 autoimmune or metabolic control traits in BRVEs (Fig. 2D). Next, we identified which brain regions were most strongly associated with a given trait. The FC-BRVEs showed the highest heritability enrichment across most traits, including depression, schizophrenia, and reaction time, which are traits previously associated with cortical brain function (Fig 2E). Some traits, including educational attainment and cognitive executive function, were also enriched in FC-BRVEs as well as hippocampus-BRVEs (Fig 2F). While others, including chronotype and morningness, were significantly enriched in both FC- and NAc-BRVEs, consistent with prior studies linking circadian rhythm to the NAc (Fig 2G)[25].

**Figure 2.**
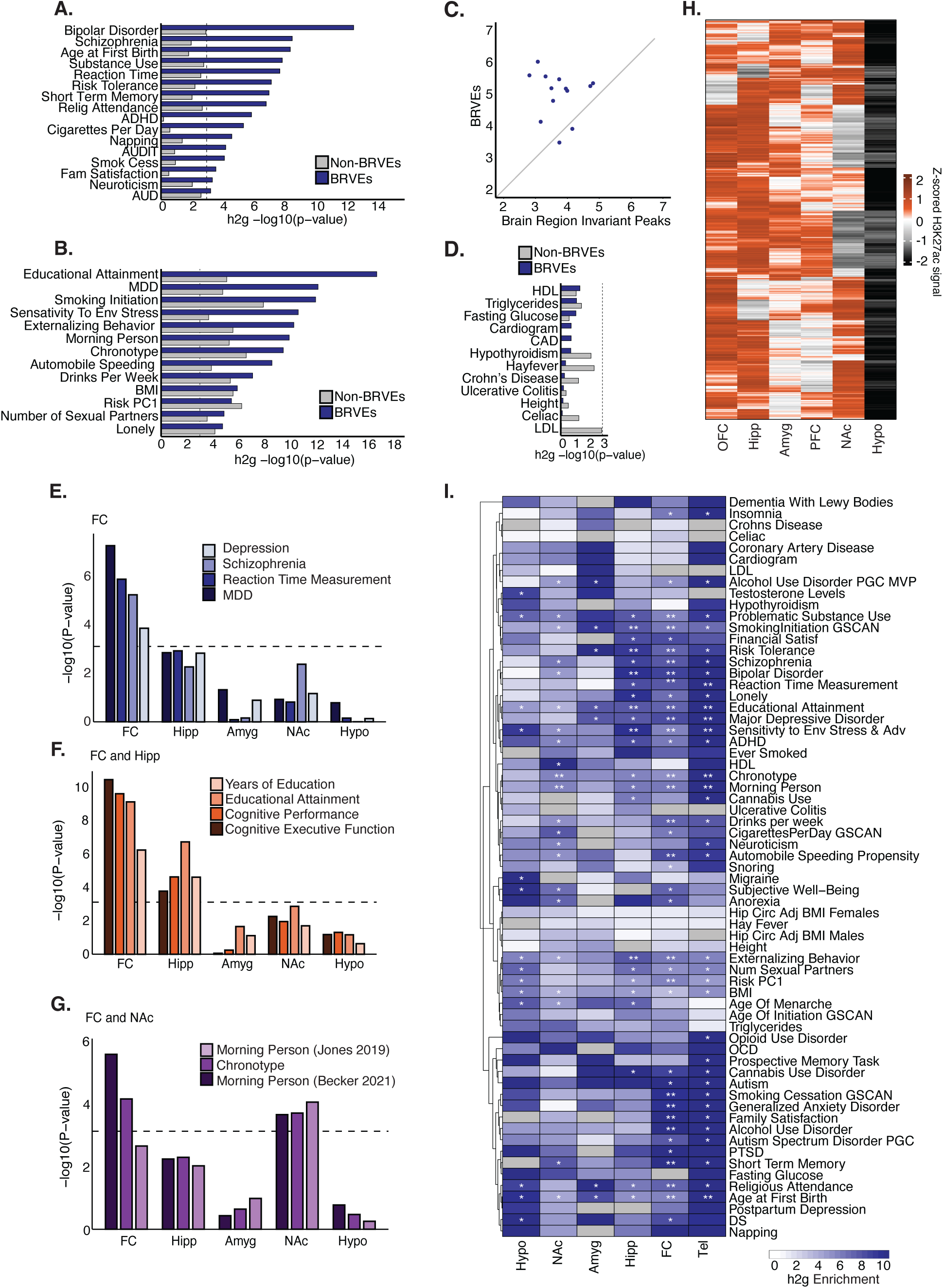
LDSC Heritability analysis of BRVEs: **(A)** Heritability enrichment analysis using s-LDSC. Shown are−log₁₀-transformed *p*-values for 16 traits that were significantly enriched in BRVEs, but not in active neuronal regulatory elements with shared activity patterns across brain regions. The dashed horizontal line indicates the multiple testing-corrected significance threshold (−log₁₀(*p*) = 3.11). **(B)** Heritability enrichment analysis using s-LDSC. Shown are −log₁₀-transformed *p*-values for 13 traits that were significantly enriched in BRVEs and regions with consistent activity patterns across brain regions. The dashed horizontal line indicates the multiple testing-corrected significance threshold (−log₁₀(*p*) = 3.11). **(C)** X-Y plot showing the enrichment value for the 13 neurological traits that were significantly enriched in both brain region-variant enhancers (BRVEs) and regions with consistent activity patterns across brain regions. **(D)** Heritability enrichment analysis using s- LDSC. Shown are −log₁₀-transformed *p*-values for 12 autoimmune or metabolic control traits. Bar plot showing heritability enrichment significance (−log₁₀(p)) for traits with strongest enrichment in the FC across brain regions (x-axis) **(E)**, traits with strongest heritability enrichment in the FC and Hipp **(F),** and traits with the strongest heritability enrichment in the FC) and NAc **(G)**. **(H)** Hierarchically clustered heatmap showing H3K27ac ChIP-seq signal across brain regions for *n* = 2,400 telencephalon-specific peaks. Values are row- normalized using z-score transformation. **(I)** LDSC heritability enrichment across 65 brain-related traits. Hierarchically clustered heatmap displaying LDSC enrichment statistics for all brain-related traits. A single asterisk (*) denotes uncorrected *p* < 0.05, and a double asterisk (**) indicates significance after Bonferroni correction (−log₁₀(*p*) = 3.11).

Of all BRVEs, 87% are associated with region-specific activity in FC, NAc, hippocampus, hypothalamus or amygdala (Fig. 1D,F). The remaining 5,337 BRVEs were not classified as specific to any one of the five regions (Fig. 1D). We explored the regional activity patterns of these remaining elements and found a striking telecephalon-specific pattern in half of these elements (n=2,400). These BRVEs show robust activity in most of the telencephalic brain regions (OFC, PFC, Hippocampus, Amygdala, and NAc) with little to no H3K27ac enrichment in the hypothalamus (Fig. 2H). These telencephalon-BRVEs had the highest enrichment values across 28 tested brain-related traits, with an average per SNP heritability enrichment above 10 (Fig. 2I). FC-BRVEs were the second highest on average across diseases with per SNP heritability enrichment of approximately 6 (Fig. 2I). Taken together, these results indicate that enhancers exhibiting diverse activity profiles across the brain are strongly linked to the heritability of neurological traits.

### DEGs linked to neuropsychiatric disorders are enriched near BRVEs

To further assess the role of BRVEs in neurological disease, we next compared BRVEs to gene expression studies of neuropsychiatric disease. We identified putative target genes of BRVEs (FDR 0.05) using a 100kb-distance-based model (methods). These enhancer-gene connection maps were highly enriched in promoter capture Hi-C and HiChIP datasets for available brain regions (Fig S2A-B). We identified 10 datasets from 8 published studies that performed differential gene expression analysis in post mortem human brain tissues for neuropsychiatric disorders, including opioid use disorder, schizophrenia, bipolar disorder and autism spectrum disorder, in brain regions profiled in our cohort including prefrontal cortex, nucleus accumbens and hippocampus [26–33]. For many of these cohorts, we found that upregulated DEGs were enriched among the full set of BRVE gene targets (Fig 3A). In contrast, the downregulated DEGs show more modest enrichment with only two out of ten cohorts enriched for expected overlaps (Fig. 3B). However, when we stratified BRVEs by their associated brain region, we found that the downregulated DEGs were frequently enriched for genes associated with region specific BRVEs for the region profiled in the DEG study (Fig. 3C-E). For example, downregulated DEGs identified in a NAc study of opioid use disorder were enriched in NAc-BRVEs (Fig. 3C), whereas downregulated DEGs identified in the frontal cortex for OUD and ASD were enriched in FC-BRVEs (Fig. 3E). This trend was most robust in NAc- and FC-BRVEs. These enrichments suggest genes linked to region-specific enhancer variation are more prone to aberrant downregulation in disease.

**Figure 3.**
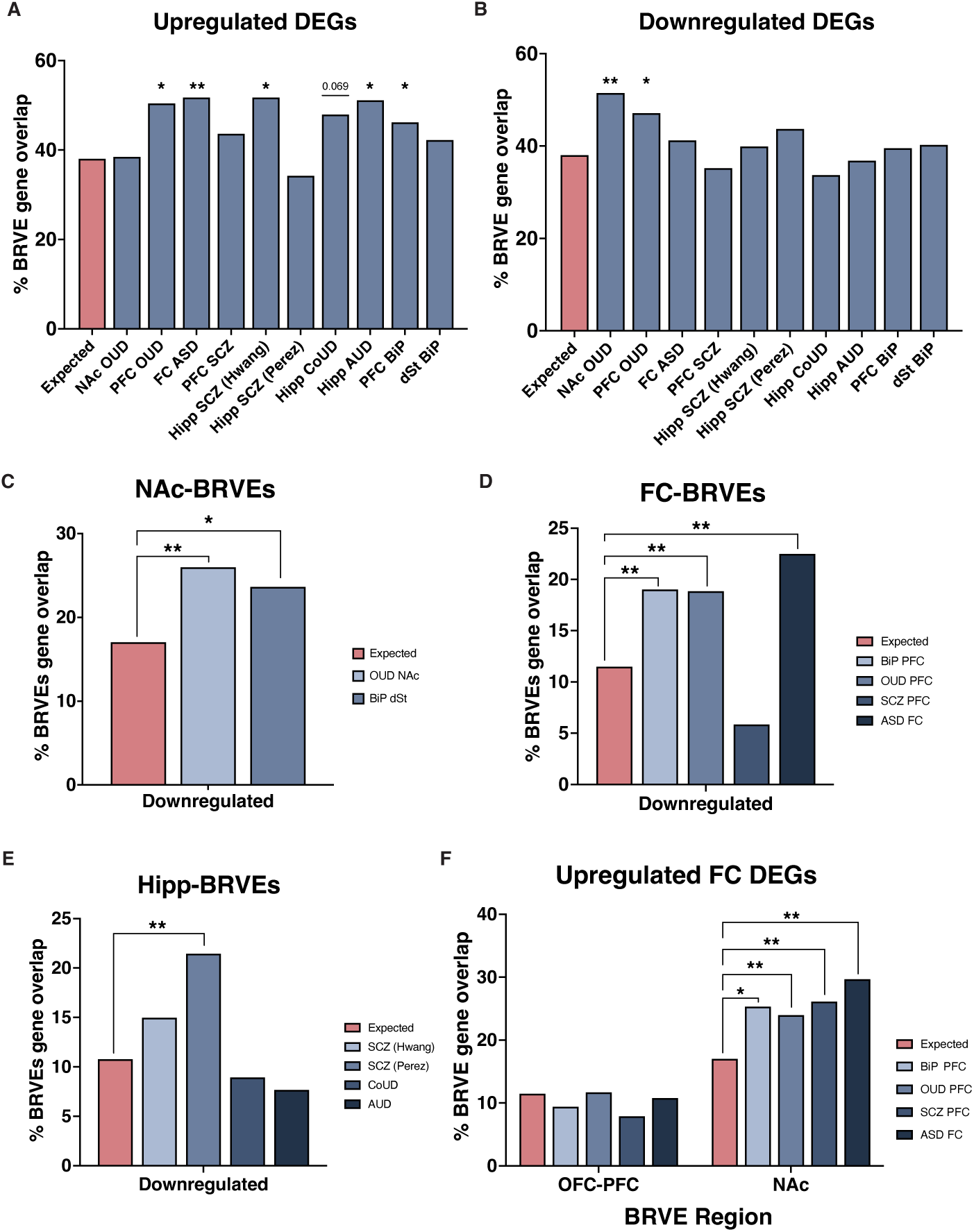
BRVE-associated genes are enriched in neuropsychiatric disorders. **(A)** Percent of upregulated DEGs identified in neuropsychiatric disorder cohorts that overlap with BRVE- associated genes. Neuropsychiatric disorders examined include opioid use disorder (OUD), autism spectrum disorder (ASD), schizophrenia (SCZ), cocaine use disorder (CoUD), alcohol use disorder (AUD), and bipolar disorder (BiP). The expected background rate is indicated by the expected bar showing the percent of all genes that overlap with BRVE-associated genes. **(B)** Same as in (A) for downregulated DEGs. **(C-E)** BRVE- associated genes for the NAc, Hipp and FC compared against DEGs downregulated in disease cohort studies from the same brain regions. **(F)** FC- and NAc-BRVE-associated genes compared against genes upregulated in the disease cohort studies of the frontal cortex. (*p-value adj < 0.05, ** p-value adj <0.001).

We further assessed the enrichment of upregulated DEGs among BRVE genes by examining region specificity. This revealed a striking pattern. Genes identified as upregulated in the FC in schizophrenia, bipolar disease, OUD and autism were consistently enriched amongst the NAc-BRVE target genes (Fig 3F). In contrast, these DEGs were not enriched in FC-BRVE target genes. This suggests that enhancers and genes typically active in the NAc may be upregulated in the FC in disease contexts. We evaluated the expression pattern of FC uprgulated DEGs associated with NAc-BRVEs in healthy brain tissue samples and found 50% of these genes to be at least moderately expressed (average TPM > 10) in the NAc across 5 donor brains. Overall, these genes were expressed approximately 1.5-fold higher in the NAc when compared against 6 PFC healthy donor brains, with 29.26% expressed at least 2-fold higher in the NAc (Fig S2C). Pathways analysis of FC upregulated DEGs linked to NAc-BRVEs revealed enrichment for BDNF signalling pathways and extracellular matrix-related terms (Fig S2C).

Collectively, these results suggest that aberrant activation of NAc-BRVEs may be a common feature of neuropsychiatric gene dysregulation.

#### Variation in enhancer activity across brain regions links ISL1 dysregulation in the NAc to BMI

Genome-wide analyses provide systems-level insights into the tissues and cell types that are broadly important for disease pathogenesis. However, these approaches do not pinpoint the particular tissue or cell type perturbed by any one disease allele. Given the strong link between disease biology and BRVEs, we reasoned that enhancer activity patterns could be leveraged to infer the cellular pathology of individual disease-associated alleles. Defining the precise cellular and tissue context for each allele is critical for interpreting its mechanistic role in disease, prioritizing relevant experimental models, and guiding the design of targeted interventions.

We compared GWAS SNPs and variants within linkage disequilibrium (LD, r^2^>0.8) to BRVEs. One of the loci with pronounced regional specificity was a BMI-associated locus near ISL1 (Insulin Gene Enhancer Protein) (Fig. 4A). Although BMI serves as a proxy for adiposity and metabolic disease risk, GWA studies have demonstrated that genetic variants associated with BMI preferentially colocalize with regulatory elements and genes active in the brain and central nervous system [34]. We observed strong clusters of BRVEs overlying this GWAS locus associated with high activity in NAc and hypothalamus, with minimal H3K27ac signal in other brain regions. In total, there were 6 NAc BRVEs and 3 Hypo BRVEs that overlapped LD SNPs at this locus. This led to the prediction that risk SNPs at this locus may act in hypothalamic neurons, NAc neurons, or both. This prediction is supported by published studies that have shown that perturbing ISL1 in hypothalamic neurons disrupts energy balance and feeding behavior, leading to obesity-related phenotypes, supporting a pathogenic role for ISL1 dysregulation in these cells [35]. We sought to determine whether ISL1 dysregulation in NAc neurons similarly contributes to BMI heritability.

**Fig. 4.**
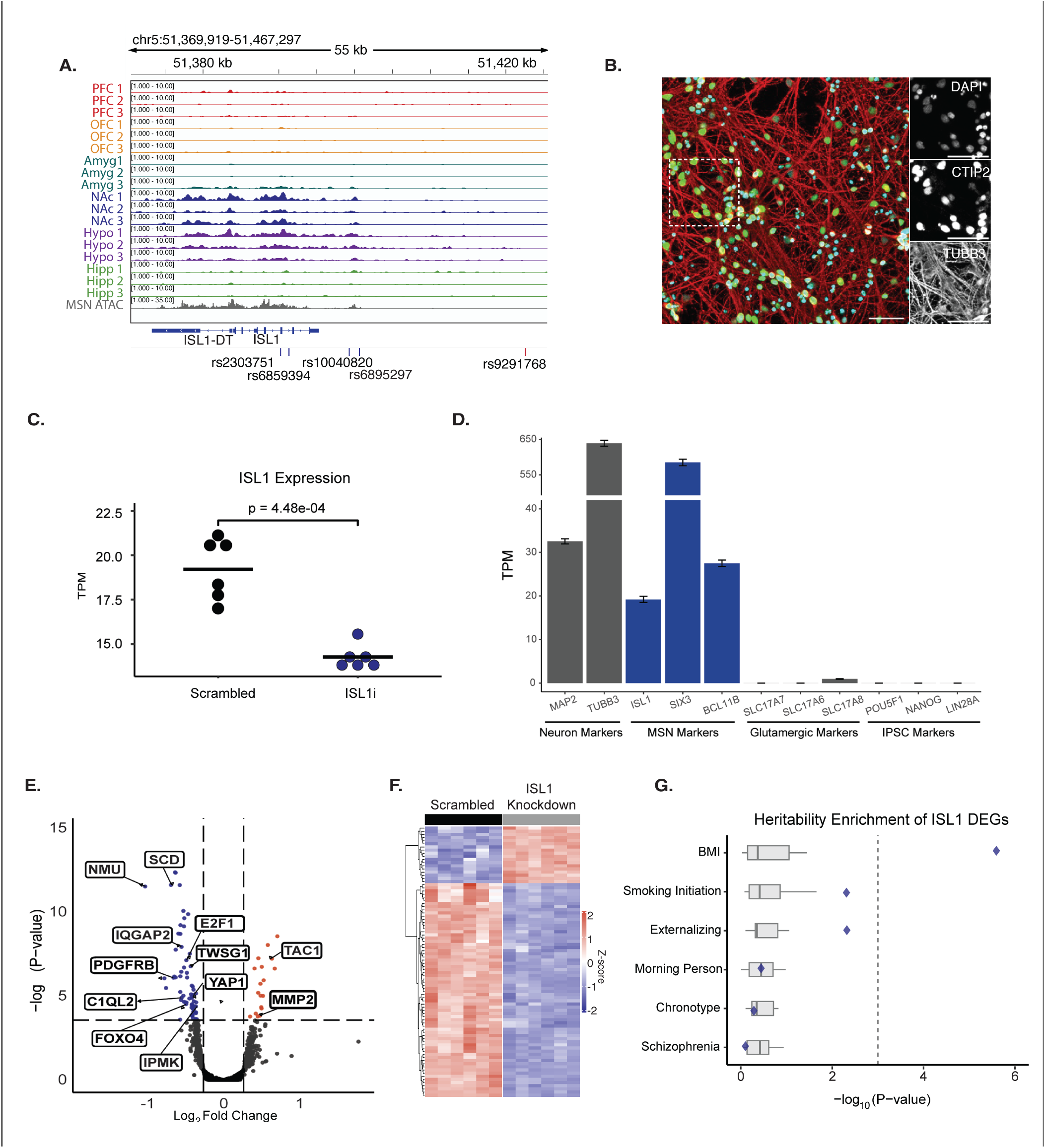
Enhancer activity variation across brain regions links ISL1 dysregulation in the NAc to BMI: **(A)** H3K27ac ChIP-seq signal across brain regions at the ISL1 locus, together with snMSN ATAC-seq signal. SNPs within the linkage disequilibrium (LD, r2 >0.8) are shown below, with the lead SNP highlighted in red and LD proxy SNPs shown in blue. **(B)** Representative image of iMSNs after 40 days *in vitro* (DIV40). Left: Composite image of DAPI, CTIP2, and TUBB3 staining in iMSNs. Scale bar = 50μm. Right: iMSNs robustly express CTIP2 (middle) and TUBB3 (bottom). **(C)** ISL1 expression is decreased in iMSNs following an ISL1 targeted siRNA knockdown (P-value 4.48E-4, T-test). (**D)** Expression of marker genes in DIV40 iMSNs. iMSNs showed high expression of genes associated with general neuronal markers and MSN-specific markers and low expression of genes associated with glutamatergic neurons and iPSCs. **(E)** Volcano plots of differential gene expression analysis of ISL1 knockdown compared to scrambled control. Upregulated DEGs are in red while downregulated DEGs are in blue (Benjamini-Hochberg adjusted P-value < 0.05 and fold-change > 1.2). **(F)** Heatmap of differentially expressed genes following ISL1 knockdown. **(G)** Heritability enrichment analysis using s-LDSC for 100 Kb-windows surrounding ISL1 knockdown DEGs. Diamonds indicate −log₁₀-transformed *p*-values for downregulated DEGs. Boxplots show −log₁₀- transformed *p*-values for 20 random subsets of n-matched gene sets selected from MSN expressed genes.

We differentiated induced pluripotent stem cells to medium spiny neurons (iMSNs) as previously described [36, 37]. We verified iMSN identity using immunohistochemistry and observed activation of marker genes TUBB3 and CTIP2 (Fig 4B). We then knocked down ISL1 in six biological replicates of iMSNs using siRNAs and observed a 1.3-fold downregulation (Fig 4C). This is aligned with modest effect sizes expected for GWAS SNPs in cis regulatory elements [38–40]. We observed a strong correlation across replicate scrambled controls and replicate ISL1 knockdowns (r^2^> 0.96), suggesting that while iPSC differentiation can yield heterogeneous populations of cells, the composition of cells is highly reproducible across replicates. We assessed the expression of iMSNs for key marker genes identified in RNA-seq studies of the brain and observed robust expression of neuronal markers, MAP2 and TUBB3 and MSN-specific gene markers, including SIX3 and BCL11B. We observed modest to low expression of markers for other neuronal subtypes such as SLC17A7 and iPSC markers like NANOG (Fig. 4D).

We identified 86 downstream differentially expressed genes associated with ISL1 knockdown (Fig 4E,F). This included genes such as NMU, which encodes a neuropeptide involved in energy homeostasis and feeding behavior [41] and IPMK, an important component of nutrient sensing and mTOR and AKT signaling pathways [42, 43]. We sought to determine whether ISL1 dysregulation in MSNs contributes pathogenically to BMI heritability. Using s-LDSC we found downstream DEGs associated with ISL1 knockdown in iMSNs were significantly enriched for BMI heritability. To assess the robustness of this enrichment, we identified n-matched control gene sets randomly selected from MSN-expressed genes that were not impacted by ISL1 knockdown and found no significant heritability enrichment in any of these 20 random sets (Fig. 4G). These results suggest that ISL1 dysregulation in MSNs is a contributor to BMI heritability and support the expansion of the prediction approach to additional cell types and loci.

### Variation in enhancer activity reveals region and cell type-specific activity of individual GWAS alleles

We next aimed to expand this approach to snATAC-seq data to predict the likely cell type of action of GWAS risk loci. GWAS loci have been shown to frequently co-localize with clusters of active enhancers arranged in cis, referred to as locus control regions, super enhancers, stretch enhancers, and multiple enhancer variant loci [3, 44, 45]. These highly cell type-specific clusters are associated with robust interconnected chromatin interactions reflecting complex enhancer-gene connectivity that is heavily pronounced at GWAS loci [46]. Moreover, GWAS loci have been shown to frequently involve multiple functional DNA variants within LD that collectively influence gene expression and genetic risk [44, 47–53]. Likewise, the ISL1 locus involved multiple BRVE peaks overlapping SNPs in LD with the GWAS SNP.

We hypothesized that the pronounced clustering of enhancers and their chromatin activity at GWAS loci could be exploited to distinguish the most likely cell type for individual disease alleles. We reasoned that the aggregate ATAC signal across all enhancers that overlap SNPs within an LD block would provide an informative metric for identifying the pathogenic cellular context of a given locus. We developed the approach, GWAS-LOCATE (GWAS LOcus Cell-type Assignment using Tissue Enhancers). The predictions described below are available for interactive exploration at https://gwas-locate.wi.mit.edu.

For each of 23,494 GWAS loci associated with neurological traits, we identified all called ATAC peaks that colocalize with the reported GWAS SNP or its LD partners (see methods) using pseudobulk signals from 42 cell clusters previously characterized across 42 brain regions from 3 donor tissues [5]. Unsupervised clustering of these ATAC signal patterns identified seven distinct groups of GWAS loci with cell type-specific activity signatures (Fig 5A,B). One of the seven clusters identified by GWAS-LOCATE, representing 24% of loci, showed diffuse activity across both neuronal and glial populations and thus these loci were deemed broadly active or indiscernible. Four clusters mapped to specific neuronal subtypes: glutamatergic cortical IT neurons (27% of loci), GABAergic MSNs (10%), and two additional GABAergic neuronal clusters: one including PVALB- and somatostatin-expressing neurons (PVALB/SST) and the other loci with specific activity in vasoactive intestinal polypeptide-expressing neurons (VIP). Glial-enriched activity accounted for 19% of loci, subdivided into microglia-specific (4%) and combined astrocyte/oligodendrocyte (15%) clusters. These activity patterns enabled us to predict the putative cell type of action for each locus. To evaluate these predictions, we linked GWAS loci to putative target genes using a distance-based model (see methods) and examined whether gene expression patterns aligned with the predicted cell-type. Genes associated with loci predicted to act in a given cell type exhibited both a higher proportion of actively expressed genes and higher mean expression levels in the predicted cell type. (Fig. 5C, Fig 5SA).

**Fig 5.**
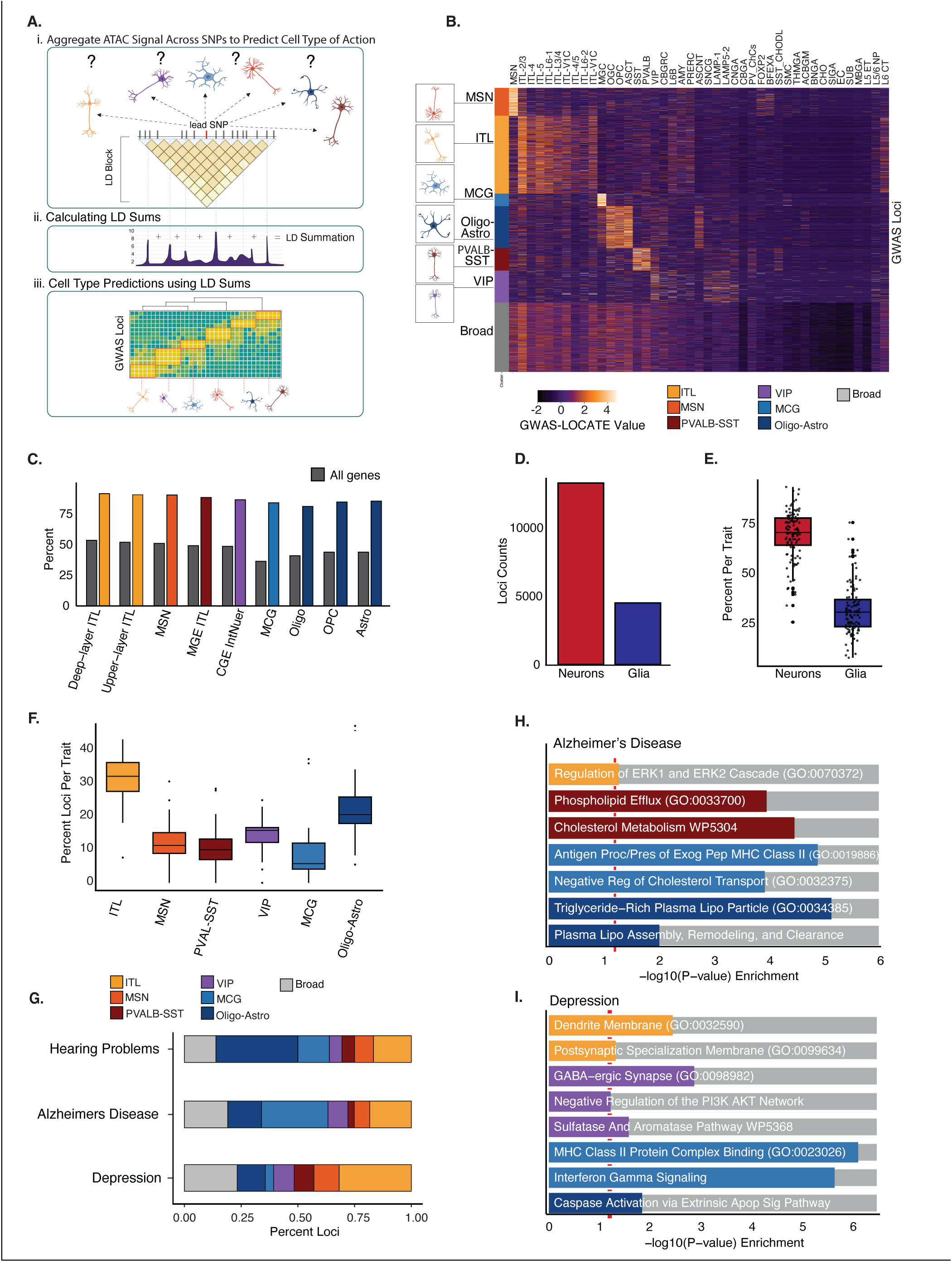
GWAS-LOCATE Clusters Reveal Cell Type Predictions of GWAS Loci: **(A)** Schematic of GWAS-LOCATE. **(i)** GWAS SNP and its LD partners are identified and compared to snATAC-seq datasets in order to predict the likely cell type of action **(ii)** ATAC peak activity accumulated GWAS SNP and LD partners. **(iii)** Summated snATAC values are clustered to identify loci with cell type specific activity patterns. **(B)** Heatmap showing the snATAC summation values for each GWAS loci. The values are row-wise z-scored. Rows are the individual GWAS loci (n = 23,494 loci) and the columns are snATAC defined cell type clusters. (left) Color bar indicates rows that correspond to GWAS loci that have similar cell type activity patterns including Oligo-Astro (dark blue), MCG (light blue) VIP (purple), PVALB/SST (dark red), MSN (orange), ITL (yellow), Broad (light gray). The columns are sorted using hierarchical clustering. **(C)** Percent of genes within 100-kb of a lead SNP predicted to act in a given cell type that are actively expressed (defined as >median RPKM) in the corresponding cell type. Percentage of all genes that are defined as “actively expressed” in each cell type is shown in grey, compared to genes associated with GWAS-LOCATE predicted loci displayed as color bars that correspond to cell type clusters shown in (B). Abbrev. Astro (astrocytes); CGE (caudal ganglionic eminence-derived interneurons); Deep-layer ITL, (deep-layer intratelencephalic neurons); Upper-layer ITL, (upper-layer intratelencephalic neurons); MSN, (medium spiny neurons); MGE, (medial ganglionic eminence-derived interneurons); MCG, (microglia); Oligo, (oligodendrocytes); OPC, (oligodendrocyte precursor cells); **(D)** Boxplots show distribution of percentages of GWAS loci predicted to act in each cell type for each trait. Each data point represents the percentage of loci predicted to act in a given cell type for an individual trait. **(E)** Bar plot showing the number of GWAS loci predicted to act in neuronal (ITL, MSN, PVALB/SST, and VIP) or glial (microglia and Oligo-Astro) cell types. **(F)** Box plot showing the percentage of GWAS loci predicted to act in neuronal or glial cell types across traits. Each data point represents the percentage of loci predicted to act in neuronal (ITL, MSN, PVALB/SST, and VIP) or glial (microglia and Oligo-Astro) cell types for an individual trait. **(G)** Stacked bar plot showing the proportion of GWAS loci predicted to act in each cell type across representative traits. Bar plots showing the significance of gene set enrichment analysis for genes associated with Alzheimer’s disease risk SNPs **(H)** and Depression **(I).**

Prior studies have shown that neuronal-specific enhancers are highly enriched for disease heritability, while glia-specific enhancers have minimal trait heritability enrichment, with Alzheimer’s disease being a notable exception [5–7]. Consistent with these results, we find the majority (75%) of neurological risk loci to be predicted to act in neurons (Fig. 5D). However, despite the paucity of genome-wide enrichment in glia-specific enhancers, we found all neurological traits to have at least one locus predicted to act in glia. 88% of traits had more than 10% of loci mapped to glial cell types (Fig. 5E).

We further evaluated the predictions by examining the percentage of loci per disease that are assigned to a given cell type. Across the five major cell group classifications, a similar number of ATAC-seq peaks were discovered (∼150,000–200,000 per group), indicating that differences in prediction rates are unlikely to be driven by variation in ATAC coverage or peak discovery. For most traits, the majority of loci linked to the trait were assigned to glutamatergic cortical IT neurons (Fig 5F). A small subset of traits has the largest fraction of loci linked to the oligodendrocyte/astrocyte cluster. This included hearing loss and menopause age (Fig. 5G, S3B). Another group of traits that includes Alzheimer’s disease (AD) and late-onset AD had a high fraction of risk loci predicted to act in microglia (Fig 5G, S3B.) Overall, the top enriched cell type from individual-locus assignments largely reflects the patterns of genome-wide heritability partitioning. These results suggest that few complex neurological traits have monolithic cellular pathology, rather the genetic risk variants linked to each trait implicate multiple cell types.

We next evaluated whether the pathways and gene sets implicated by distinct cell-type predictions revealed distinct biological pathways that underpin complex traits. For each trait, we linked all loci predicted to act in the same cellular context to genes using a 100-kb distance model and performed gene ontology and pathway enrichment analysis. For Alzheimer’s disease, we predicted 29% of loci to act in microglia. Amongst the microglia predicted loci, we find a significant enrichment for genes involved in cholesterol metabolism and MHC class II activity. In contrast, AD loci predicted to act in cortical neurons were enriched for genes involved in ERK1/2 regulation. This pattern is consistent with prior work showing that cholesterol handling and antigen presentation are core microglial functions in AD [54]. ERK1/2 dysregulation has been linked to tau phosphorylation, AB plaque formation and neuroinflammation, implicating both neuronal and microglia roles for ERK1/2 in AD pathology [55] (Fig 5H). For depression, we found 9% of GWAS loci were predicted to act in VIP neurons and the associated genes were enriched for GABA receptor activity. This finding is consistent with GABAergic deficit models of depression, in which altered GABA receptor signaling and interneuron dysfunction contribute to the pathology [56]. Depression loci predicted to act in the microglia were enriched for genes involved in interferon gamma signaling. Interferon gamma has been shown to prime microglia, leading to depression associated behaviors in model organisms [57] (Fig 5I).

Collectively, these results suggest that localizing GWAS risk loci to their predicted cell type of action via GWAS-LOCATE can reveal distinct shards of the pathology of complex traits. While some of these pathways may be discovered by evaluating gene set enrichment across all loci without regard to cell type of action, others such as ERK1/2 dysregulation in cortical neurons for Alzheimer’s and GABA-ergic synapse in VIP neurons for depression are not identified when all loci are combined prior to pathway analysis, suggesting grouping all loci without regard to cell type may obfuscate disease-critical pathways that are driven by loci that act in minor cell types.

## Discussion

Our understanding of the epigenetic landscape of the human brain has drastically expanded in recent years. First by improved protocols to perform ChIP-seq in post mortem brain tissues [58, 59] and later by the expansion of single cell epigenetic techniques. However, there are still comparatively few unique donors that comprise reference datasets and some brain regions remain to be fully characterized. Here, we incorporate sorted neuron and glial datasets from six brain regions in biological triplicate to build on these important reference maps. Consistent with prior studies, we find that the majority of variation in enhancer and gene activity is linked to major differences in cell type, with the greatest axis of variation in the brain being the distinction between neurons and glial cell lineages. Our study demonstrates that while enhancers with variation across the brain represent the minority of elements, these elements play a disproportionate role in disease biology. Whether this variation in these elements is fully driven by cell composition such as MSN in the NAc or reflect variation in activity of similar neuronal subtypes in different brain regions, both suggest the need to continue to expand epigenetic datasets to better capture this variation.

We found BRVEs to be disproportionately enriched for genetic variation linked to neurological traits and genes that are differentially expressed in post mortem brain studies of human disease. This is consistent with the growing body of literature demonstrating that complex traits are primarily driven through cell type-specific rather than essential or ubiquitous genes. These results are consistent with a model in which complex neurological traits are driven through hundreds of small effect size genetic variants, and suggest these variants may be maintained in the population due to their limited context-specific effects [60–63]. These results highlight the value in profiling DEGs in multiple disease-relevant brain regions to further understanding of trait biology.

They also demonstrate the need to perform functional dissection of risk loci in the appropriate cellular context. As an example we highlighted the ISL1 locus. ISL1 is a transcription factor that is expressed in multiple cellular contexts, thus association alone is insufficient to determine the effector cell type. One of the ongoing challenges to the study of enhancer biology is linking enhancers with the correct target gene. Accurately determining the target gene of a particular enhancer region is highly dependent on cell type and cellular context. Recent advances in Hi-C and its deviations, functional approaches such as CRISPR inhibition and computational prediction strategies [64], have helped identify target genes of cis acting regulatory elements. For each of these strategies, however, the cell type or cell model utilized has a major impact on the identified target gene. Thus, identifying the appropriate pathogenic cell type is critical to identifying the disease-relevant gene target.

Our approach, GWAS-LOCATE, aims to reveal the critical cell type that is impacted by a given disease locus. While genome-wide analyses can reveal the dominant, often termed “causal” cell type for a given disease, the individual variants or loci that contribute to disease biology are unlikely to uniformly impact a single cell type. Dissecting trait biology by cell type can reveal novel disease insights by enabling correct mapping of enhancer to target genes or by revealing gene sets or patterns that are indistinguishable when hundreds of risk loci are combined for gene network or pathway analyses. Knowing the pathogenic cell type of a risk locus also has important implications for therapeutic development as treatment delivery strategies may be cell type dependent. We provide predictions for 23,494 loci through integration with snATAC-seq study. One limitation of this approach is the 24% of loci we identified as “indiscernible” or broadly active. This could indicate loci that broadly impact multiple cell types in the brain or that the most appropriate cell type or context is not included in this analysis. As our approach relies on cell type specific ATAC patterns there are inherent limitations in cell type resolution, particularly for loci that may act in many cell types. Future studies that incorporate different developmental stages, environmental exposures, additional brain regions or disease-specific enhancer profiling may help further distinguish cell type for these loci.

**Supplemental Table 1**: Brain Tissue Samples

**Supplemental Table 2:** Brain Region Variable Elements (BRVE) coordinates

**Supplemental Table 3:** Traits in heritability analysis

**Supplemental Table 4:** Differential gene expression analysis of ISL1 knockdown

## Methods

### Sample acquisition and ChIP-sequencing

Tissue samples (n = 18) from the dorsolateral prefrontal cortex, orbitofrontal cortex, hippocampus, amygdala, nucleus accumbens, and hypothalamus were collected were obtained from cadaver donors collected by the University of Miami Brain Endowment Bank and the NIH NeuroBioBank These samples were anonymized and collected after death, thus their use does not constitute human subjects research and is therefore exempt from regulation 45 CFR Part 46 (NIH SF424 Part II: Human Subjects). Nuclei were sorted using NeuN (Catalog number: MAB377X, vendor: Sigma Aldrich, manufacturer: Chemicon) as a marker of neuronal nuclei followed by ChIP-seq for H3K27ac Catalog number: 39133, vendor: Active Motif) as previously described [65].

### ChIP-sequencing data processing and normalization

Adapter sequences were removed from paired-end reads using Cutadapt v1.9.1. Reads shorter than 20bp were removed [66]. Filtered reads were aligned to the hg38 genome assembly using BWA-MEM v0.7.17-r1188 in paired-end mode with default parameters [66, 67]. Output SAM files were converted to BAM format and indexed using SAMtools v1.10. Peaks were identified with MACS v2.1.2 [68, 69]. The bigWig files were evaluated using Integrative Genomics Viewer and samples exhibiting low signal-to-noise were eliminated from the study [70]. Quality control metrics were generated using the ChIPQC Bioconductor package, and libraries with very low mapping rates, RelativeCC enrichment <1, reads in peaks <2%, or fewer than 10,000 peaks were excluded [71]. BigWig tracks were normalized to reads per kilobase per million (RPKM) with 50 bp bins and visualized on Integrative Genomics Viewer (IGV). The bigWig tracks of each region were aggregated through merging via bigWigMerge, bedSort, bedGraphToBigWig. All called peaks were filtered to remove ENCODE blacklisted regions [72].

Inherent normalization was used to quantify H3K27ac ChIP reads as a part of the Axiotl Platform as previously described [73]. This approach uses positive control genomic regions, defined from Roadmap 2015 reference datasets to identify regions with stable epigenomic activity across cell types and tissues. Negative control regions are defined as peaks with activity in only one reference panel cell type. The average signal across positive and negative controls is compared to “inherently” normalize ChIP signal. This method is detailed in prior studies [73, 74]. All browser track images are adapted from IGV.

### Identification of Brain Region Variable Elements (BRVEs)

Variation in H3K27ac ChIP-seq signal across brain regions was assessed using one-way analysis of variance (ANOVA) implemented with the scipy.stats.f_oneway function in SciPy [75]. Six groups corresponding to brain regions were included in the analysis, each consisting of three biological replicate values per group. P-values obtained from the one-way ANOVA performed for each peak were adjusted for multiple testing using the Benjamini–Hochberg false discovery rate (FDR) procedure, as implemented in statsmodels.stats.multitest.fdrcorrection in Python [76] [77]. The FDR threshold was set at α = 0.10. 40,049 peaks passed the Benjamini– Hochberg significance criterion and were retained for downstream analysis.

### Classification of BRVEs by brain region specificity patterns

Following the one-way ANOVA, post hoc analyses were performed using independent two-tailed t-tests implemented in Python with scipy.stats.ttest_ind [75]. For each peak, the triplicate samples corresponding to a given brain region were compared against the combined samples from the remaining five brain regions (each-versus-all else comparison). This procedure was performed iteratively for each brain region and for each peak in the dataset.

A p-value was obtained for each brain region per peak. Fold changes shown in 1F reflect the mean signal of the focal brain region compared to the mean signal across the remaining brain regions (each-versus-all else comparison). Peaks with p<0.05 in these post-hoc tests were used to define BRVEs for each brain region.

### Comparison of BRVEs to snATAC

Published snATAC-seq data obtained from were compared to BRVEs [5]. snATAC-seq peak annotations from pseudobulk analysis from 42 cell clusters were compared to BRVE peaks. The percentage of BRVEs overlapping active ATAC peaks were identified.

#### Mapping BRVEs and SNPs to putative target genes

Candidate peak-to-gene interactions were assigned using two methods: 100-kb window approach or the nearest gene approach. For the 100-kb window method, BRVEs were assigned to all genes with transcriptional start sites within 100-kb of peak center and defined as BRVE-associated genes. GWAS loci evaluated using GWAS-LOCATE were likewise assigned to all genes within 100-kb of the lead GWAS SNP coordinate.

### Heritability enrichment using stratified Linkage Disequilibrium Score Regression

Linkage disequilibrium score regression (LDSC) was used to estimate heritability enrichment for brain-related traits within genomic regions of interest. Regions of interest included, SNPs within 2.5kb of BRVE peak centers and SNPs within 100-kb of DEGs identified with ISL1 knockdown. LD scores for annotated SNPs were calculated using European ancestry reference panels from HapMap3 and the 1000 Genomes Project Phase 3, consistent with the ancestry of the GWAS cohorts, following the previously described approach [78, 79].

GWAS summary statistics for relevant traits were obtained from the NHGRI-EBI GWAS Catalog [80]. To standardize file formats and ensure proper harmonization across studies, summary statistics were processed using the Bioconductor package MungeSumstats [81]. The baseline LDSC model described by Bulik-Sullivan et al. was used to regress GWAS test statistics on LD scores for annotated SNPs to estimate partitioned heritability. Heritability enrichment was calculated by comparing the proportion of total SNP heritability attributable to variants within the defined regions to the proportion of genome-wide SNPs they represent.

### Comparison of BRVE-associated genes with disease gene expression studies

Genes associated with BRVEs were compared to previously published post mortem disease studies [26–33]. Disease studies included: opioid use disorder data sets extracted from the prefrontal cortex, NAc, and midbrain; autism spectrum disorder data sets extracted from the frontal cortex; schizophrenia data sets extracted from the prefrontal cortex and hippocampus; cocaine use disorder data set extracted from the hippocampus; alcohol use disorder data set extracted from the hippocampus; and bipolar disorder data sets extracted from the prefrontal cortex and dorsal striatum. Enrichments were evaluated by comparing the percent of DEGs that are associated with BRVEs vs. the percent of all genes that are associated with BRVEs (using 100-kb distance approach) for each region and dataset.

### GWAS-LOCATE: using snATAC-seq to predict cell type of action for individual GWAS associations

An interactive browser to explore the results of GWAS-LOCATE,described below, is available at: https://gwas-locate.wi.mit.edu. This website also allows custom downloads of disease or cell type specific subsets of predictions, as well as access to the full set of predictions.

Lead SNPs associated with neurological traits were obtained from the NHGRI-EBI GWAS Catalog[80]. Variants in LD with each lead SNP were identified using SNIPA with an r² threshold of 0.8 [82].We used published pseudobulk snATAC-seq peak calls, in which accessible chromatin regions were identified using MACS2 and represented as fixed 501-bp peaks [5]. In the original study, a cell-by-bin count matrix was generated for each sample, merged across samples, and converted to a binary accessibility matrix to indicate the presence or absence of accessible chromatin at each region. For our downstream analyses, each published 500-bp peak was expanded to a fixed 1,500-bp window centered on the peak midpoint to standardize the genomic intervals. The snATAC peaks were then intersected with the lead SNPs and their LD partners. For each reported GWAS SNP, we compared the lead SNP and its LD partners to the binary matrix of snATAC-seq peaks to calculate the total sum of active peaks that coincide with putative causal variants associated with each GWAS locus. We refer to this as the “GWAS-LOCATE score”. We calculated the GWAS-LOCATE scores for each of the 42 cell clusters, using pseudobulk binary peak calls. This resulted in a GWAS-LOCATE score for each of the 42 cell clusters and each reported GWAS SNP, represented as a matrix in Figure 5B. GWAS-LOCATE scores for each lead SNP across all cell types is available at: URL: https://gwas-locate.wi.mit.edu.

Assigning LD Sum snATAC Loci to Putative Cell type of Action

We clustered the matrix of GWAS-LOCATE scores to identify subsets of GWAS SNPs that had shared ATAC-seq cell type specificity patterns using K-means clustering (scikit-learn, KMeans, n_clusters = 7, random_state = 0). Seven clusters were selected for k-means clustering based on a scree (elbow) analysis evaluating cluster numbers ranging from 0 to 20. The elbow at 7 clusters indicated a diminishing marginal gain in explained variance with additional clusters, supporting the choice of seven as an optimal balance between model complexity and information capture.

Gene Ontology and Pathway Enrichment

Gene Ontology and pathway enrichment analyses were performed using Enrichr.

Python and R Environments

Python v.3.8.10 and R v.4.2.1 were used.

## Generation of iMSNs

Induced pluripotent stem cell-derived medium spiny neurons (iMSNs) were differentiated from a female human iPSC line that constitutively expresses dCas9-KRAB-ZIM3 marked by mCherry. This clonal cell line was derived from the Whitehead Institute’s Stem Cell facility from fibroblasts obtained from a healthy donor at Coriell Cell Repository. Prior to differentiation, iPSCs were maintained on Matrigel-coated plates in mTeSR Plus media. All media are changed daily throughout the differentiation protocol. Upon ∼80% confluency, iPSCs are dissociated into single cells using accutase, passaged into Matrigel-coated 6-well plates at a density of 4x10^5^ cells per well in mTeSR Plus media, and expanded for 1 day. Then, for the first 5 days *in vitro* (DIV), basal media (composed of DMEM/F12, N2 supplement, B27 without retinoic acid, and GlutaMAX supplement) with dual SMAD inhibition (using SB431542 and LDN193189) is applied to the cells. On DIV5, cells are dissociated into single cells using accutase and passaged into Matrigel-coated plates in basal media with dual SMAD inhibition, sonic hedgehog (SHH), and Wnt signaling pathway inhibition (using DKK-1) at a density of 3x10^6^ cells per well. Then, between DIV12 and DIV21, basal media with SHH and DKK-1 is applied. Cells are passaged using accutase into PLO/Laminin-coated 6-well plates at a density of approximately 1x10^6^ cells per well on DIV21. During DIV21 through DIV25, cells are exposed to basal media with B27 with retinoic acid, SHH, DKK-1, and brain-derived neurotrophic factor (BDNF). Starting on DIV26 and continuing on until the end of experiments, cells are maintained in maturation media, which is composed of basal media, B27 with retinoic acid, and BDNF. All experiments are conducted after DIV30.

## Immunofluorescence staining of iMSNs

On DIV 40, cells were fixed with cold paraformaldehyde for 20 minutes (min) at room temperature (RT). Following three washes in phosphate-buffered saline with calcium and magnesium (PBS+/+), cells were permeabilized for 10 minutes at RT in 0.1% Triton X-100. Then, cells were blocked in PBS+/+ with 10% donkey serum for 1 hour at RT. Cells were incubated overnight in primary antibodies diluted 1:100 in PBS+/+ containing 1% donkey serum at 4°C. After overnight incubation, cells underwent three 20-minute washes with PBS+/+, then incubated for 90 minutes in secondary antibody (1:1000 dilution) with PBS+/+ and 1% donkey serum at RT in the dark. Following three PBS+/+ washes, nuclei were counterstained with DAPI for 2 minutes. Fixed cells were then washed with PBS+/+ and subsequently imaged. Images were acquired using a ZEISS LSM 980 confocal microscope with a 20× objective. Image analysis was performed using Fiji (ImageJ, NIH).

## siRNA Knockdown and bulk RNAseq

iMSNs (n=6 biological replicates) in 6-well plates were exposed to siRNAs targeting ISL1 on DIV 37 for 72h. Cells were washed with 1mL phosphate-buffered saline (PBS), then incubated with 500μL OptiMEM I Reduced Serum Medium and 300μL of transfection mixture containing Lipofectamine RNAiMAX, Opti-MEM I Reduced Serum Medium, and siRNA of choice for 1 hour at 37°C. Scrambled siRNA sequence functioned as the negative control. After 1 hour, 3mL of iMSN maturation media was applied. Cells were then left undisturbed until collection timepoint on DIV40. During collection, cells underwent a 1mL PBS wash followed by cell scraping and collection in 700μL of QIAzol Lysis Reagent, then stored in microcentrifuge tubes at -80°C until RNA extraction with Qiagen’s miRNeasy Mini Kit. Extracted RNA was then submitted for bulk-RNA sequencing (paired end, 150x150bp, NOVASEQS1, prepared with Watchmaker-mRNA kit).

## ISL1 knockdown and differential gene expression analysis

RNA-seq read quality was assessed using FastQC (v0.11.9)[83]. Fastp (v0.23.4) was then used to trim the reads to exclude low-quality sequences and sequencing adapters[84]. Utilizing STAR (v 2.7.1a)[85], the trimmed reads were mapped to the human hg38 genome and the GENCODE annotated transcriptome (release V46). featureCounts (v2.0.1)[86] was applied to count the number of reads mapped to each gene. Only genes with CPM > 3 in at least six samples were retained and normalized using the TMM method[87]. Differentially expressed genes were identified using edgeR’s (v3.40.2)[88] likelihood ratio-based test with a nonzero fold-change built into the null and an FDR cutoff < 0.05 (Supplementary Table 4). and Oligo-Astro.

## Acknowledgements

This study was funded by DP1DA044337.

## Data Access and Availability

An interactive browser to explore these predictions described below is available at: https://gwas-locate.wi.mit.edu/ . This website also allows custom downloads of disease or cell type specific subsets of predictions, as well as access to full set of predictions.

ChIP-seq tracks and BRVE calls are also available https://gwas-locate.wi.mit.edu/ .

## Author Contributions

OC, HK, YS, KH conceived the project and designed the experiments.

YS, KH, ZB, CA performed laboratory experiments.

HR, YS CO, AH, GM, CA and OC performed computational analysis.

HR, YS, KH, CA, CO, and MM prepared figures and visual displays of data.

HR, YS, KH, CO, ZB, AH, WD, YH, XL, CA, and OC provided intellectual input on data interpretation and analysis.

DM prepared and disseminated brain tissue.

HR, YS, CO and OC wrote the manuscript with input from all co-authors.

---

**Supplemental Figure 1.**
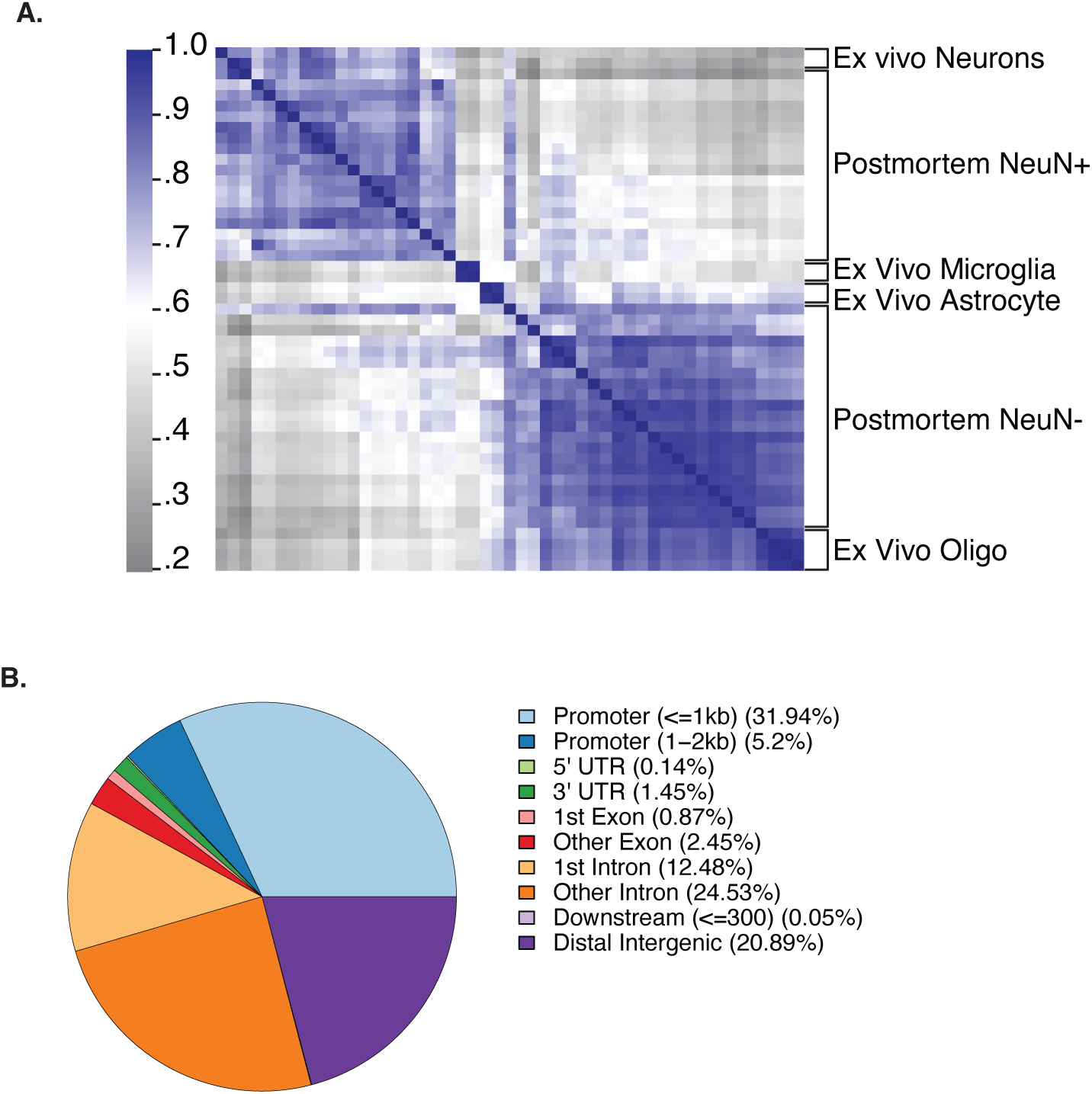
: Comparative analysis of H3K27ac ChIP-seq from six brain regions: **(A)** Correlation clustermap showing Pearson correlation coefficients from normalized ChIP-seq signal values. The clustermap compares ex vivo neurons, ex vivo microglia, ex vivo astrocytes, and NeuN⁺ and NeuN⁻ postmortem ChIP-seq samples across six brain regions. **(B)** Genomic annotation of all brain H3K27ac peaks (FDR < 0.10) generated using ChIPseeker. Peaks were annotated relative to hg38 gene features (TxDb.Hsapiens.UCSC.hg38.knownGene) with promoter regions defined as ±2 kb of the transcription start site (TSS). Promoter-associated peaks were further subdivided into those located within **1 kb** of the TSS (Promoter ≤1 kb) and those located **1–2 kb** from the TSS (Promoter 1–2 kb). The pie chart shows the proportion of peaks assigned to promoter, untranslated region (UTR), exon, intron, downstream, and distal intergenic genomic features.

**Supplemental Figure 2.**
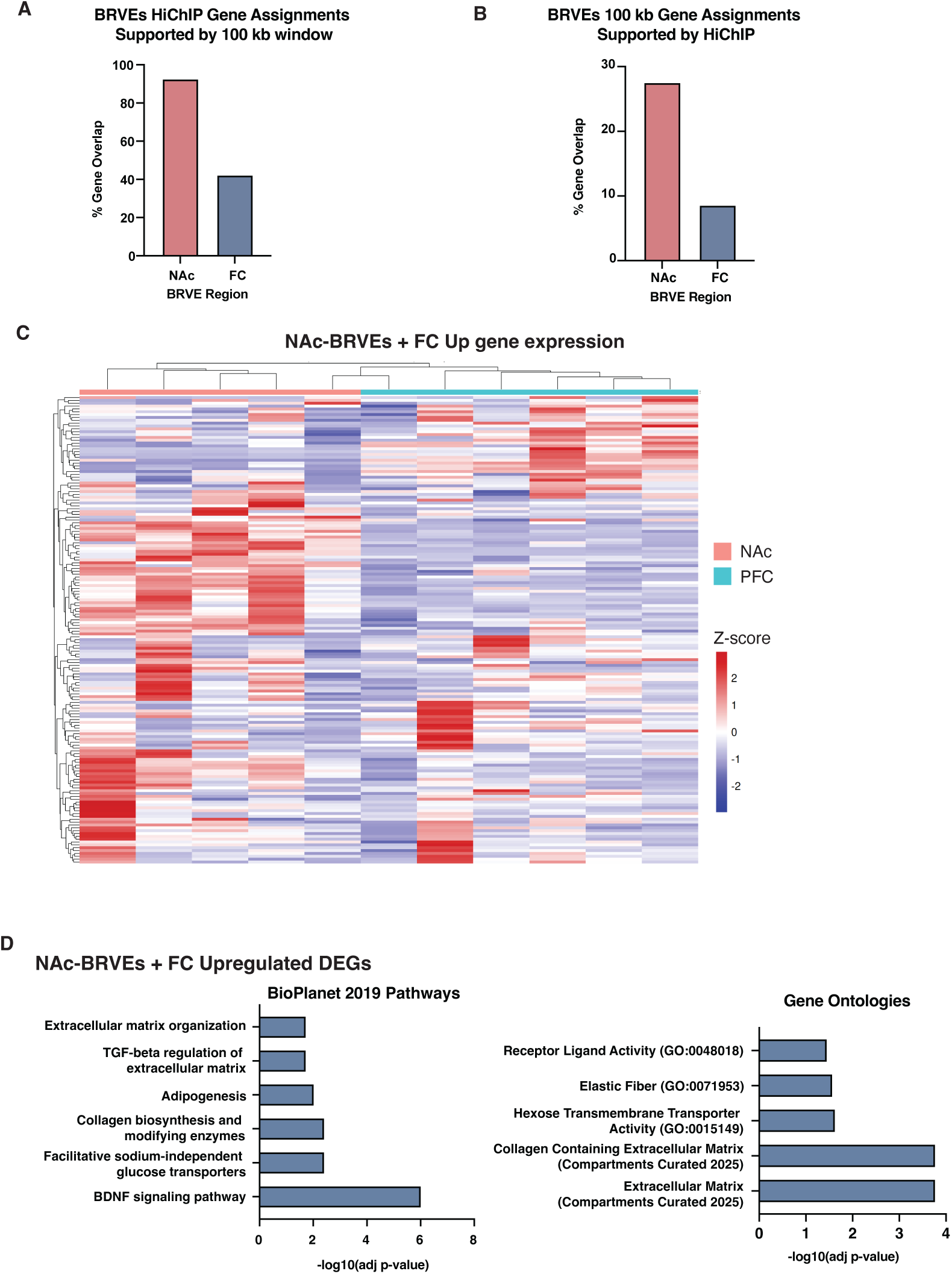
: Gene expression analysis of neuropsychiatric disease genes: **(A)** Percent of NAc- and FC-BRVEs assigned by HiChIP that are supported by the ±100 kb window approach of peak-to-gene mapping. **(B)** Percent of NAc- and FC-BRVEs assigned by the ±100 kb window approach that are supported by HiChIP peak-to-gene mapping. **(C)** Gene expression in healthy NAc (n=5) and PFC (n=6) tissues. Genes shown are associated with NAc-BRVEs and upregulated in the frontal cortex in neuropsychiatric disease. Heatmap is z-scored by row with zeros removed. **(D)** BioPlanet 2019 Pathways and Gene Ontology analyses of genes that are upregulated in the frontal cortex in neuropsychiatric disorders and defined as NAc-BRVEs through the ±100 kb window approach.

**Supplemental Figure 3.**
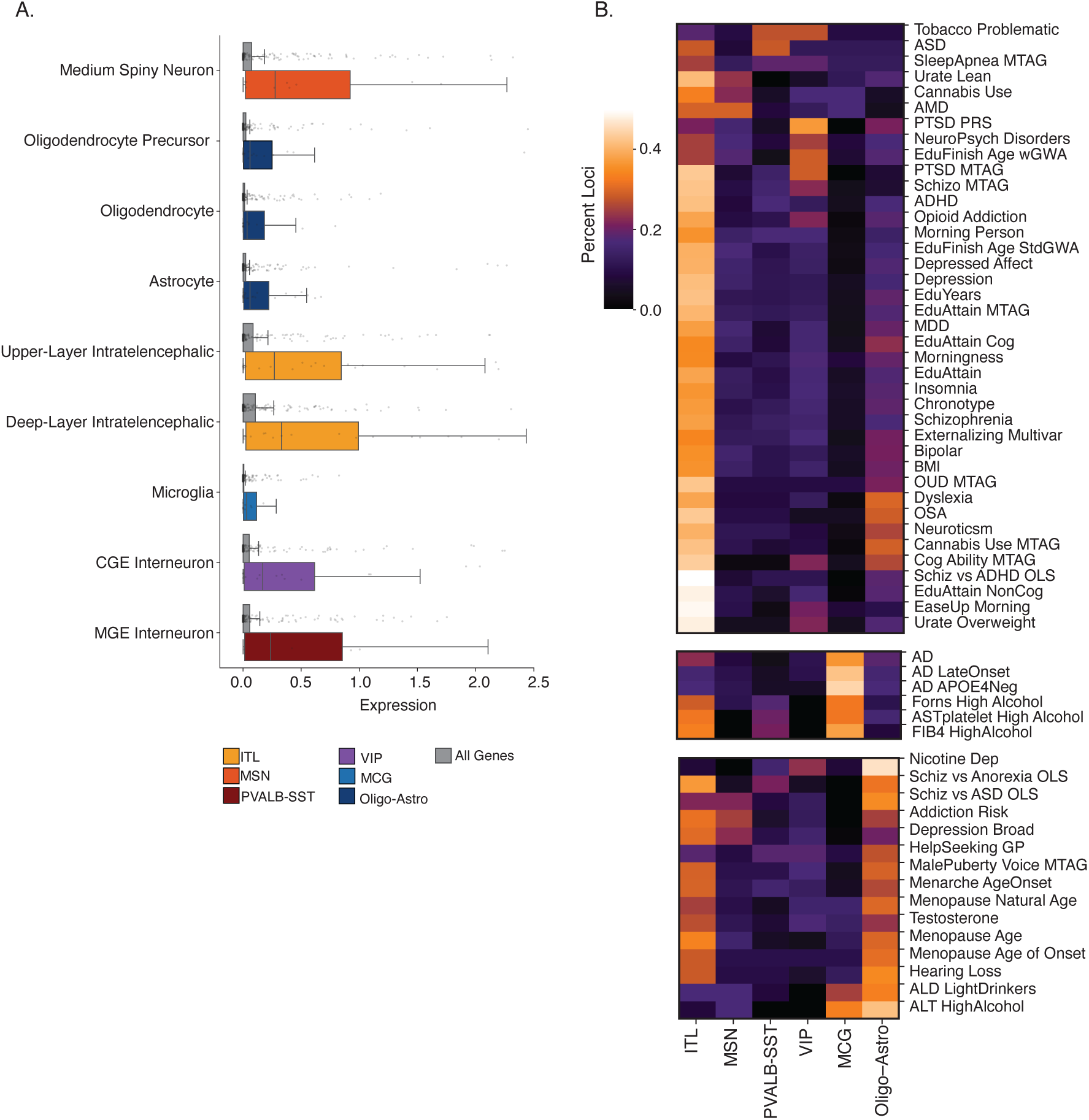
: GWAS-LOCATE cell type proportions and putative target gene expression: **(A)** Distribution of gene expression levels of all genes that are defined as “actively expressed” in each cell type is shown in grey, compared to genes associated with GWAS-LOCATE predicted loci displayed as color bars that correspond to cell type clusters from Fig5B. . Abbrev. Astro (astrocytes); CGE (caudal ganglionic eminence-derived interneurons); Deep-layer ITL, (deep-layer intratelencephalic neurons); Upper-layer ITL, (upper-layer intratelencephalic neurons); MSN, (medium spiny neurons); MGE, (medial ganglionic eminence-derived interneurons); MCG, (microglia); Oligo, (oligodendrocytes); OPC, (oligodendrocyte precursor cells); **(B)** Heatmap showing the percentage of loci predicted to act in each cell type for selected traits. Traits are grouped according to the cell type with the highest proportion of predicted loci, including Intratelencephalic (ITL) (top), Microglia (MCG), and Oligo-Astro.

